# Neural circuit mechanisms of sexual receptivity in *Drosophila* females

**DOI:** 10.1101/2020.08.07.241919

**Authors:** Kaiyu Wang, Fei Wang, Nora Forknall, Tansy Yang, Christopher Patrick, Ruchi Parekh, Barry J. Dickson

**Affiliations:** Janelia Research Campus, Howard Hughes Medical Institute, 19700 Helix Drive, Ashburn VA 20147, U.S.A; Queensland Brain Institute, University of Queensland, St Lucia QLD 4072, Australia

## Abstract

Choosing a mate is one of the most consequential decisions a female will make during her lifetime. This is particularly true for species in which females either mate repeatedly with the same partner or mate infrequently but use the sperm from a single copulation to fertilize eggs over an extended period of time. *Drosophila melanogaster* uses the latter strategy. Here, we characterize the neural circuitry that implements mating decisions in the female brain. A female fly signals her mating choice by opening her vaginal plates to allow a courting male to copulate^1,2^. Vaginal plate opening (VPO) occurs in response to the male courtship song and is dependent upon the female's mating status. We sought to understand how these exteroceptive (song) and interoceptive (mating status) inputs are integrated to control VPO. We show that VPO is triggered by a pair of female-specific descending neurons, the vpoDNs. The vpoDNs receive excitatory input from vpoEN auditory neurons, which are tuned to specific features of the *melanogaster* song. The song responses of vpoDNs, but not vpoENs, are attenuated upon mating, accounting for the reduced receptivity of mated females. This modulation is mediated by pC1 neurons, which encode the female’s mating status^3,4^ and also provide excitatory input to vpoDNs. The vpoDNs thus directly integrate the external and internal signals to control the mating decisions of *Drosophila* females.

---

*Drosophila* males woo their potential mates by vibrating their wings to produce a species-specific courtship song. This song comprises two main components: brief trains of high-amplitude pulses (pulse song) and continuous low-amplitude oscillations (sine song)^5,6^. The male song induces deflections of the female aristae, thereby activating auditory sensory neurons (JONs) that project to the central brain^7^. Several types of song-responsive neurons have been identified in the female brain^8–11^, but it is unknown if and how these neurons regulate sexual receptivity. How a female responds to a male’s song is largely determined by whether or not she has previously mated. Once mated, females store sperm for days to weeks, and during this time are reluctant to mate again^12^. A male seminal fluid peptide (sex peptide, SP) binds to sperm and signals the presence of sperm in the female reproductive tract through an ascending pathway from the SP sensory neurons (SPSNs) in the uterus via the SAG neurons in the ventral nerve cord to the pC1 neurons in the brain^3,4,13–15^. SP silences neuronal activity in this pathway, such that the SPSN, SAG and pC1 neurons are all less active in mated females than in virgins^4,15^, lowering the female’s sexual receptivity^3,4,13–15^. How are these distinct external and internal signals are integrated in the female brain to control VPO (Video 1), the motor output that signals a willingness to mate?

Female receptivity is impaired by blocking the activity of the ~2000 neurons that express in either of the two sex-determination genes, *fruitless* (*fru*)^16,17^ or *doublesex* (*dsx*)^3,18^. This class of neurons includes the *fru*^+^ *dsx*^+^ SPSNs^13,14^, the *dsx*^+^ SAGs^15^, and the *dsx*^+^ pC1 cells^3^. To search for other *fru*^+^ or *dsx*^+^ neurons that contribute to female receptivity, we screened a collection of 234 sparse driver lines specific for various *fru*^+^ or *dsx*^+^ cell types generated using the split-GAL4 technique^19,20^. Virgin females in which a neuronal silencer^21^ was expressed by one of these lines were assayed for their frequency of copulation within 10 minutes of being individually paired with a naive wild-type male (Fig. 1a). Of the seven lines with the strongest reduction in receptivity, two labelled the SPSNs, one the SAGs, and one the pC1 cells. The other three lines all targeted a pair of female-specific descending neurons which we named the vpoDNs (Fig. 1b; Extended Data Fig. 1a). These neurons are *dsx*^+^, *fru*^−^ and cholinergic (Extended Data Figs. 1b, 2). Their dendrites arborize primarily in the lateral protocerebrum and their axons project to multiple regions of the ventral nerve cord, including the abdominal ganglion (Fig. 1b).

**Fig 1.**
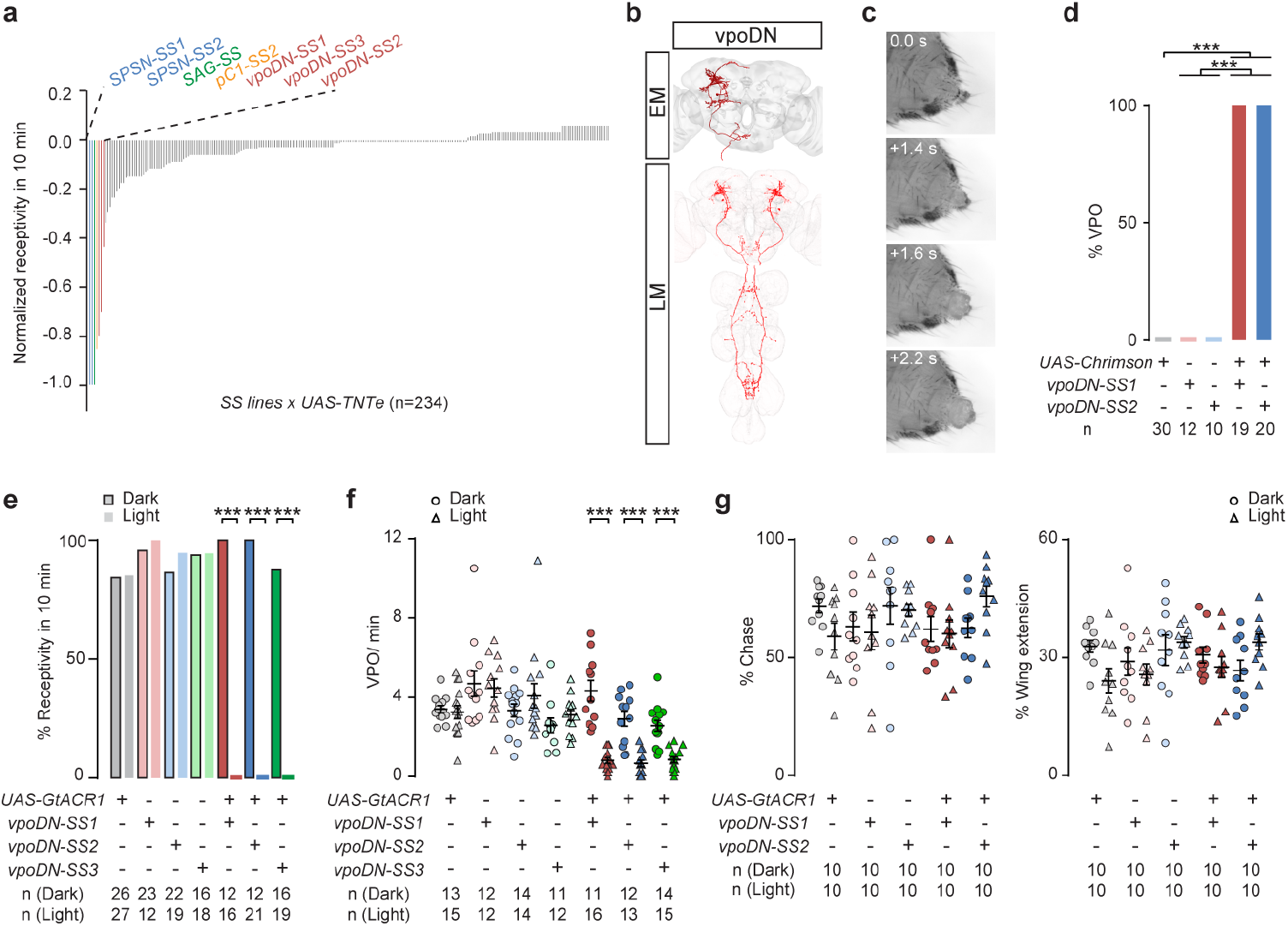
Female-specific vpoDNs control vaginal plate opening (VPO). **a**, Percentage of virgin females copulating within a 10-min observation period, for each split-GAL4 line tested (n = 234 lines, 17-36 flies per line). Results were normalized to the control group with UAS-TNTe but no split-Gal4. **b**, EM and confocal (LM) images of vpoDNs in the female central nervous system. **c**, Snapshots of female VPO induced upon photoactivation of vpoDNs (see also Video 2). **d**, Percentage of virgins exhibiting VPO in response to photoactivation. **e**, **f**, Percentage of virgins copulating (**e**) and frequency of VPO (**f**) within 10 min of being paired with a wild-type male. **g**, Percentage of time wild-type males chased or extended a wing towards a virgin female of the indicate genotype during a 10-min observation period. ***, *P* < 0.001, Fisher’s exact test in **d**and **e**, Wilcoxon test in **f**and **g**. Data in **f**and **g**shown as scatter plots with mean ± s.e.m..

The vpoDNs function as command-type^22^ neurons for VPO. Photoactivation^23^ of vpoDNs in virgin females reliably triggered VPO in isolated females (Fig. 1c, d, and Video 2). Conversely, in virgin females paired with wild-type males, acute optogenetic silencing^24^ or genetic ablation^25,26^ of the vpoDNs prevented copulation and dramatically reduced the frequency of VPO (Fig. 1e, f, Extended Data Fig. 1c, d), without diminishing the intensity of male courtship (Fig. 1g). The vpoDNs are morphologically similar to the female-specific *dsx*^+^ pMN2 descending neurons, which were proposed to function in oviposition^27^. Because the pMN2 neurons were identified by stochastic labelling, we cannot conclusively determine whether vpoDNs are indeed the same neurons as pMN2 neurons. However, we found no evidence that vpoDNs contribute to oviposition.

In addition to extending a wing to sing, males perform several other actions while courting females, including extending their proboscis, licking and holding the female, and curling their abdomen^1^. We annotated the occurrence of each of these male actions in courtship videos and found that wing extension is the most frequent of these actions just prior to VPO (Fig. 2a). We confirmed previous reports^10,28^ that copulation rates are reduced if males are muted by removing their wings or females are deafened by removing their aristae (Fig. 2b). These amputations also reduced the frequency of VPO (Fig. 2c). These results suggested that the vpoDNs might be activated by male song. Indeed, in two-photon imaging experiments using the calcium indicator GCaMP6s^29^, we detected a robust elevation of calcium levels in the soma and neurites of vpoDNs in virgin females upon playback of male courtship song (Fig. 2d). This calcium response was generally preserved, though variably diminished, when one of the aristae was glued to prevent its vibration, but completely eliminated when the second arista was also immobilized (Fig. 2d).

**Fig 2.**
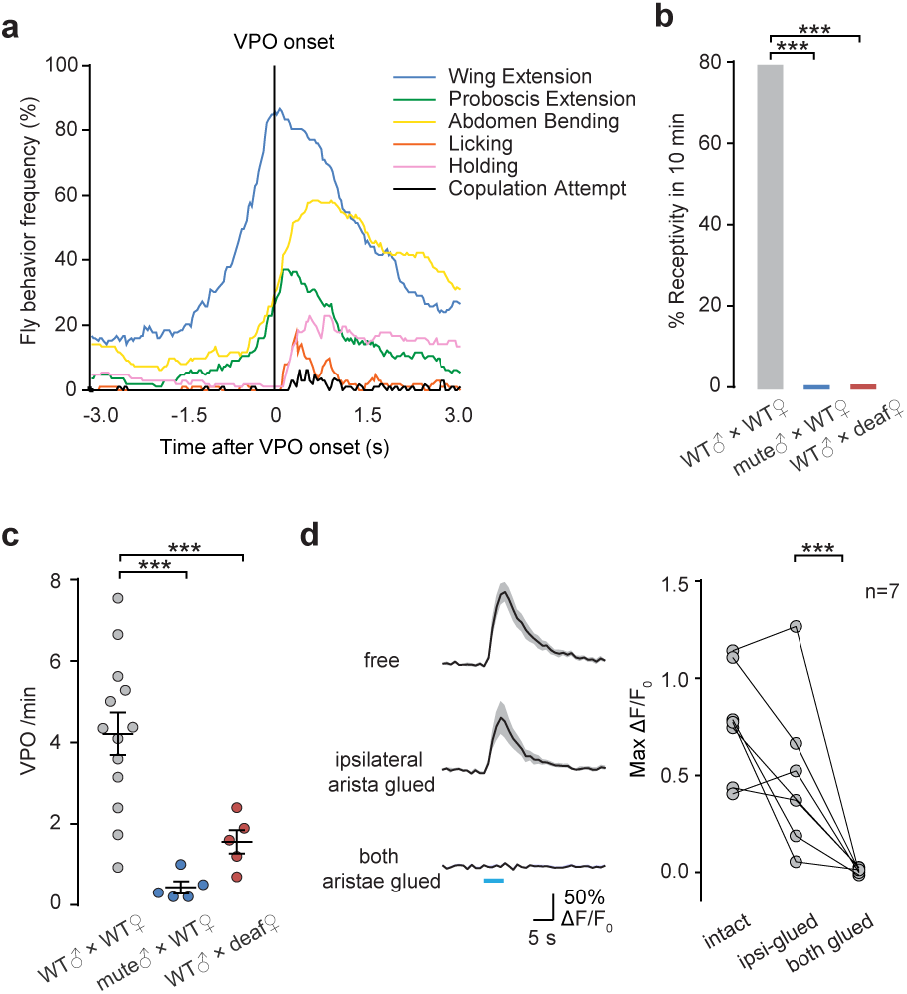
vpoDNs are sensitive to courtship song. **a**, Frequency of indicated male courtship behaviours around the onsets of female VPO, during a 10-min observation period. n = 124 VPOs from 12 pairs of flies. **b**, **c**, Percentage of virgins copulating (**b**) and frequency of VPO (**c**) within a 10-min observation period. **d**, GCaMP6s signal changes in vpoDNs in response to conspecific courtship song, with or without aristae immobilized. ***, *P* < 0.001, Fisher’s exact test in **b**, Wilcoxon test in **c**and **d**. Data in **c**and **d**shown as scatter plots with mean ± s.e.m..

The vpoDN dendrites lie mostly in the superior lateral protocerebrum with no obvious dendritic arborizations within the antennal mechanosensory center (AMMC), the primary auditory neuropil, nor of the wedge region, a secondary auditory neuropil known to include song-responsive neurons^9,10^ (Fig. 1b). We therefore sought to trace potential pathways from these auditory centers to the vpoDNs within the FAFB electron microscopic volume of the female brain^30^. We identified a single vpoDN in each hemisphere. We extensively traced the vpoDN in the right hemisphere (Fig. 1b) as well as its presynaptic partners of this right vpoDN, identifying a total of 45 neurons with at least 10 synapses onto vpoDN (Fig. 3a, Extended Data Table 1).

**Fig 3.**
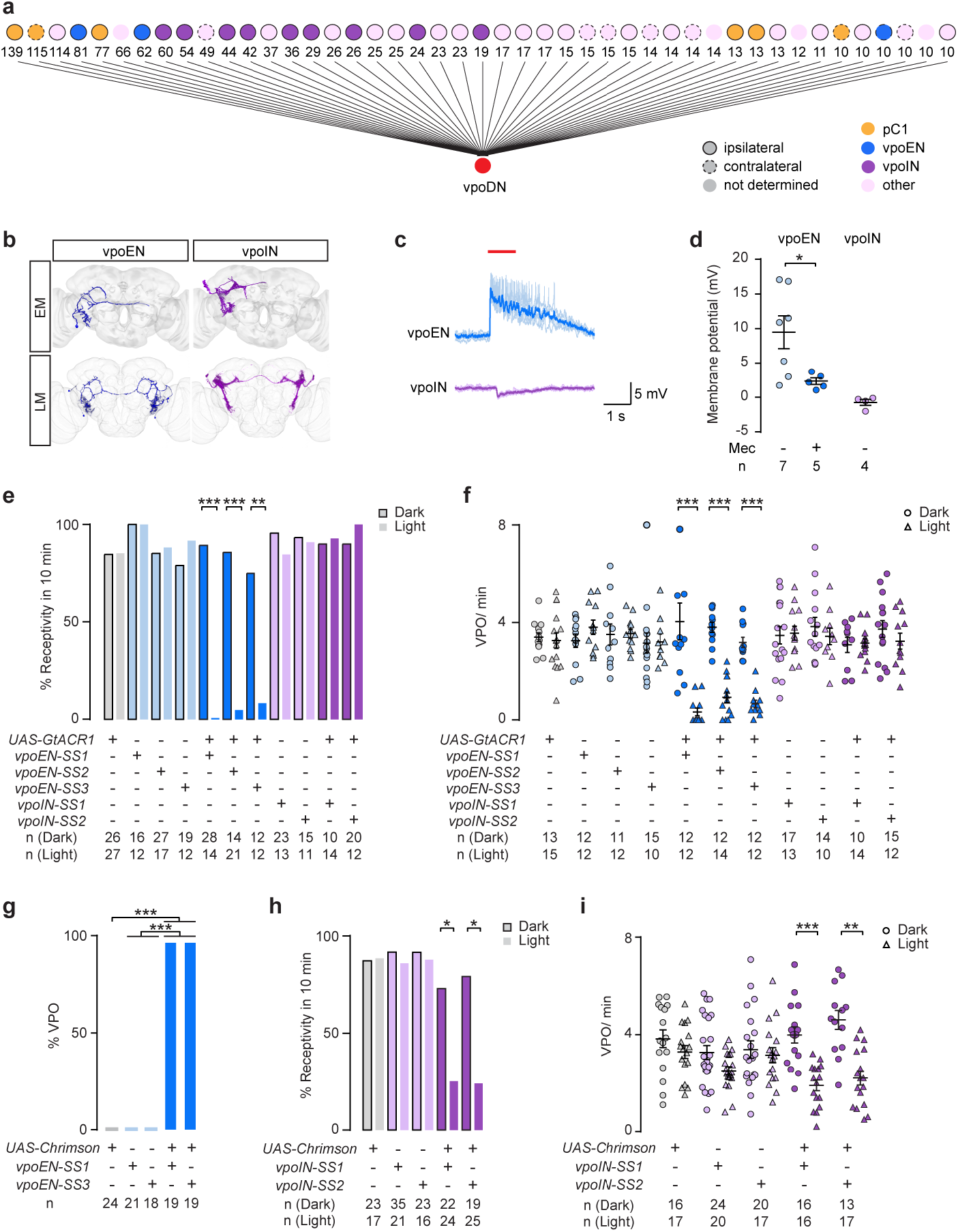
Auditory inputs to vpoDNs. **a**, Neurons presynaptic to a single vpoDN in the FAFB EM volume, showing the number of input synapses identified (thresholded at 10). **b**, EM reconstructions (top) and confocal (LM) images (bottom) of vpoEN and vpoIN cells. **c**, Example traces of membrane potential changes in vpoDNs upon photoactivation (red bar) of vpoENs or vpoINs. **d**, Plots of vpoDN membrane potential changes upon photoactivation of vpoENs or vpoINs, before and after mecamylamine (Mec) application. **e**, **f**, Percentage copulating (**e**) and frequency of VPO (**f**) for virgins of the indicated genotypes within a 10-min period of courtship by a wild-type male. **g**, Percentage of virgins exhibiting VPO in response to vpoEN photoactivation. **h**, **i**, Percentage copulating (**h**) and frequency of VPO (**i**) for virgins of the indicated genotypes within a 10-min period of courtship by a wild-type male. ***, *P* < 0.001, **, *P* < 0.01, *, *P* < 0.05, Fisher’s exact test in **e**, **g and h**, Wilcoxon test in **d**, **f** and **i**. Data in **d**, **f** and **i** shown as scatter plots with mean ± s.e.m..

None of the vpoDN input neurons received direct input from the AMMC, but at least two cell types had extensive arborizations within the wedge (Fig. 3b). We obtained multiple split-GAL4 driver lines specific for these two cells (Extended Data Fig. 3). Using these lines to label the cells for light microscopy (Fig. 3b and Extended Data Fig. 2), we determined that one of these cell types is cholinergic, and hence presumably excitatory, whereas the other is GABAergic and therefore most likely inhibitory. Accordingly, we named these two cell types the vpoENs and vpoINs, respectively (Fig. 3b). The vpoINs are morphologically similar to the previously described *fru*^+^ aSP8^31^, aSP-k ^32^ and vPN1^33^ neurons; the vpoENs do not resemble any previously described cell type. Within FAFB there are two vpoEN cells and 14 vpoIN cells in each hemisphere. We performed whole-cell recordings on the vpoDNs while optogenetically stimulating either the vpoENs or vpoINs. As expected, activation of vpoENs depolarized vpoDNs, whereas activation of vpoINs weakly hyperpolarized vpoDNs (Fig. 3c, d).

Optogenetically silencing and activating the vpoENs and vpoINs revealed that they promote and inhibit, respectively, VPO and receptivity. In assays in which virgin females were paired for 10 minutes with wild-type males, acutely inhibiting the vpoENs significantly reduced the frequency of copulation (Fig. 3e) and VPO (Fig. 3f). Conversely, in isolated females, strong optogenetic activation of vpoENs elicited VPO (Fig. 3g), mimicking activation of vpoDNs (Fig. 1d). In virgin females paired with males, activating vpoINs suppressed mating (Fig. 3h) and VPO (Fig. 3i), whereas silencing vpoINs had no obvious effect (Fig. 3e, f).

Using two-photon calcium imaging, we found that both vpoENs and vpoINs, like vpoDNs (Fig. 2d), responded to playback of male courtship songs (Fig. 4a, b). To determine whether vpoDNs, vpoENs, and vpoINs are tuned to specific features of courtship song, we generated artificial songs consisting of only the pulse or sine components. The critical song feature for species recognition and female receptivity is the interval between successive pulses within a pulse train^5,34^. In *melanogaster*, this inter-pulse interval (IPI) is approximately 35ms. For the artificial pulse songs, we therefore systematically varied the IPI between 10ms and 300ms. We also tested responses to white noise and to a distinct sound that males produce by flapping their wings during aggressive interactions^35^. Both vpoENs and vpoDNs responded robustly only to courtship pulse songs with an IPI near 35ms (Fig. 4b). Neither cell type responded to sine song, even if its amplitude was increased to match that of the pulse song, nor to white noise or antagonistic sounds (Fig. 4b). The vpoINs were much more broadly tuned, responding to pulse song across a wide range of IPIs, to the aggressive sound, and also weakly to sine song and white noise (Fig. 4b). The combination of strong excitation from highly-selective vpoENs and weak inhibition from broadly-responsive vpoINs may ensure that the vpoDNs are finely tuned to the conspecific courtship song.

**Fig 4.**
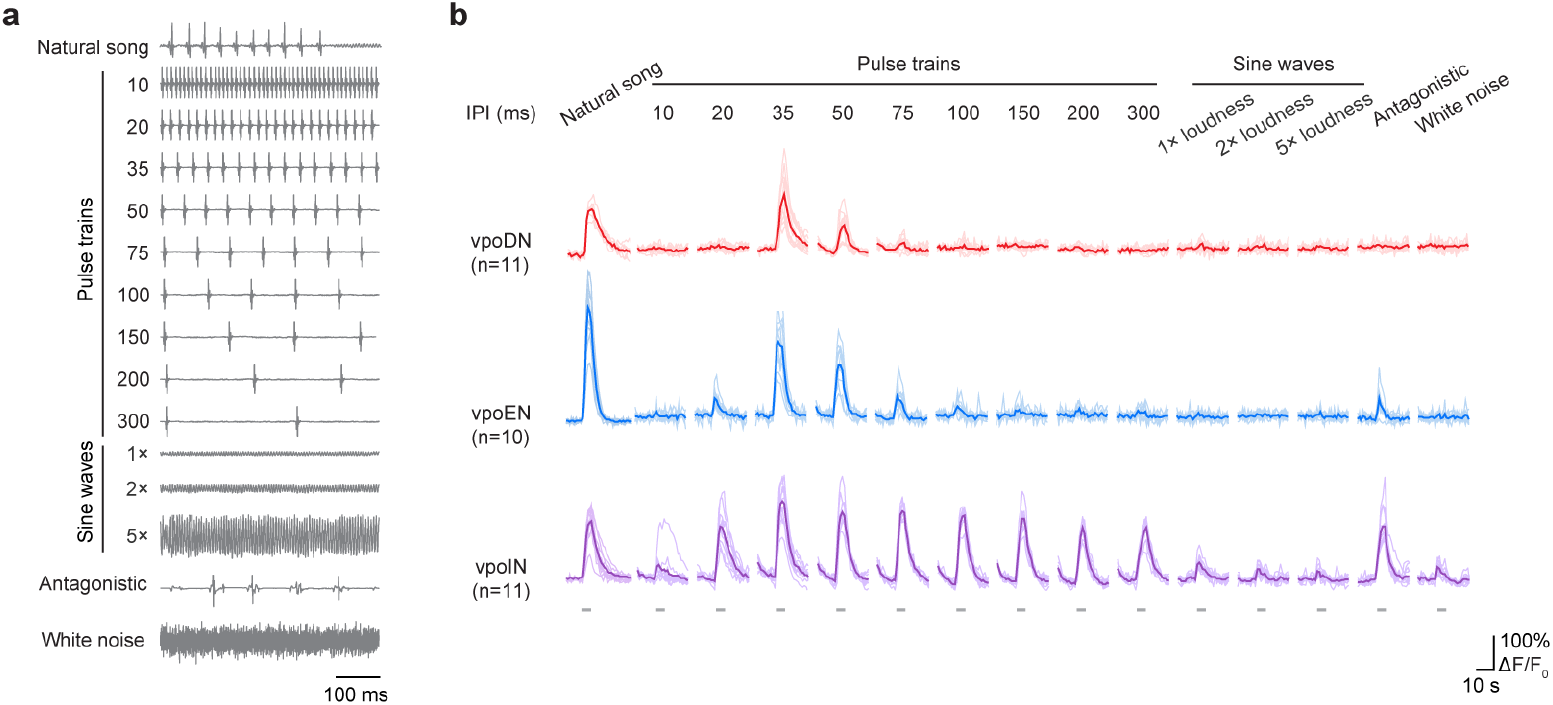
vpoENs and vpoDNs are selectively tuned to conspecific courtship song. **a**, Traces of the songs played to the virgin females. **b**, Response traces summarizing the sound-evoked GCaMP6s responses in three cell types. Darker traces were averaged from lighter ones which represent the response in different samples. Gray bars indicate stimuli (5 s).

Finally, we sought to determine how the auditory control of sexual receptivity is modulated according to the mating status of the female. Mated females are less receptive than virgins, which we found to correlate with a lack of VPO (Fig. 5a). The absence of VPO after mating suggests that the vpoENs and vpoDNs are either less potent at eliciting VPO, or are less excitable, in mated females compared to virgins. We tested these possibilities in optogenetic activation and calcium imaging experiments. We found that vpoDNs are equally potent in mated and virgin females (Fig. 5b), but that both the basal calcium levels (Fig. 5c) and the response to courtship song (Fig. 5d) were lower in mated females than in virgins. The vpoENs, in contrast, were significantly less potent at eliciting VPO in mated than in virgin females (Fig. 5b), but neither the basal activity nor song response of vpoENs were lower (Fig. 5c, d). We conclude from these data that in mated females, vpoENs are less able to excite vpoDNs upon detection of conspecific pulse song. We also imaged calcium levels in vpoINs and found that both the basal activity and song responses of these cells were indistinguishable between mated and virgin females (Fig. 5c, d). Thus, mating status information is integrated into the VPO control circuit at the level of the vpoDNs, not via either of their vpoEN or vpoIN auditory inputs.

**Fig 5.**
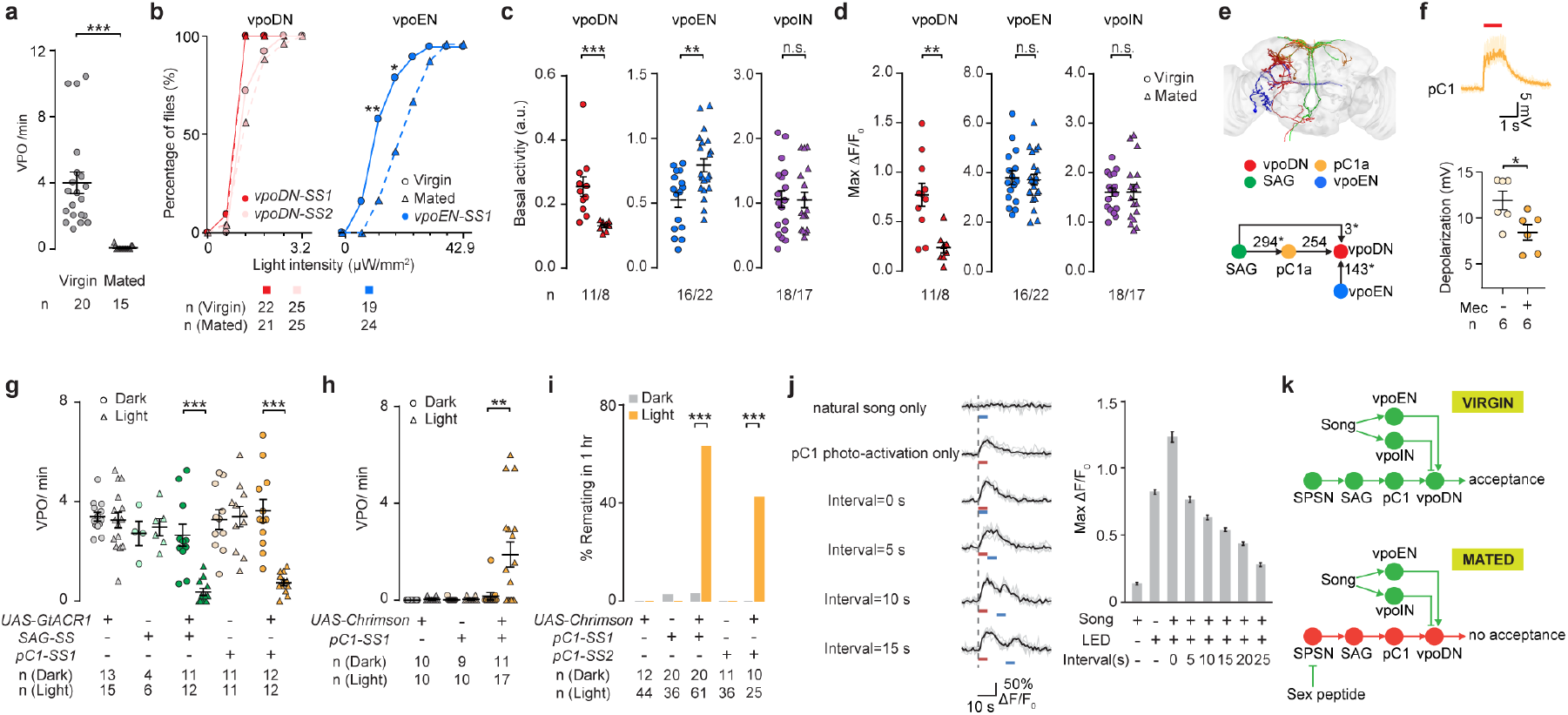
vpoDNs integrate mating status and song. **a**, Frequency of VPO for wild-type females during a 10-min courtship assay. **b**, Percentage of flies exhibiting VPO upon photoactivation of vpoDNs or vpoENs at varying light intensities. **c**, **d**, Basal (**c**) and song-evoked (**d**) GCaMP6s signals. **e**, EM reconstructions of and synapse counts among indicated cells. *, incomplete reconstruction. **f**, Example traces and plots of membrane potential changes in vpoDNs upon photoactivation of pC1 neurons (red bar) before and after mecamylamine (Mec) application. **g**, VPO frequency by virgin females during courtship with or without optogenetic inhibition of SAG or pC1 neurons. **h**, **i**, Frequency of VPO (**i**) and percentage of mated females copulating (**j**) within 1 hr of courtship with wild-type males, with or without photoactivation of pC1 neurons. **j**, Example GCaMP6s traces (left) and plots (right) of song responses in vpoDNs of mated females at various intervals after photoactivation of pC1 neurons (n = 7). **k**, Model for the integration of song responses and mating status in vpoDNs. vpoENs relay conspecific song signal to vpoDNs. In virgins, activity of the SPSN-SAG-pC1 pathway potentiates vpoDNs to facilitate the transformation of song signals into VPO. After mating, SP silences the SPSN-SAG-pC1 pathway, suppressing song responses in vpoDNs and hence female receptivity. ***, *P* < 0.001, **, *P* < 0.01, *, *P* < 0.05, Fisher’s exact test in **b** and **i**, Wilcoxon test in **a**, **c**, **d**, **f**, **g** and **h**. Data in **a**, **c**, **d**, **f**, **g** and **h** shown as scatter plots with mean ± s.e.m..

Of all the vpoDN input cells we identified in the FAFB EM volume, the cell type with the most synaptic connections was the pC1 cells (Fig. 3a, 5e, Extended Data Table 1). By performing whole-cell recordings, we found that photoactivation of pC1 neurons elicited a strong depolarization and action potentials in vpoDNs (Fig. 5f). The pC1 cells receive input from the SPSN-SAG pathway^4^. Upon mating, male SP silences this pathway, so that all of these neurons have lower activity in mated females than in virgins^4,15^. We hypothesized that mating suppresses vpoDN excitability at least in part by silencing the pC1 cells, a major excitatory input to vpoDNs. In support of this hypothesis, we found that acutely silencing either SAG or pC1 neurons in virgin females suppressed VPO to the low frequency normally observed in mated females (Fig. 5g). Conversely, photoactivation of pC1 cells in mated females restored both VPO and sexual receptivity (Fig. 5h, i). Most importantly, we found that transient (5 seconds) photoactivation of the pC1 neurons in mated females rendered the vpoDNs more sensitive to courtship song, directly demonstrating that pC1 cells control vpoDN excitability. Interestingly, we also noted that this effect persisted for up to 25 seconds after photoactivation of pC1s (Fig. 5j).

In summary, our data reveal the cellular and circuit basis for sexual receptivity in *Drosophila* females. The decision to mate or not to mate is largely determined by how the vpoDNs integrate signals from two direct synaptic inputs: the vpoENs, which are selectively tuned to the conspecific male courtship song, and the pC1 cells, which encode the female’s mating status (Fig. 5k). When the male sings, female vpoENs would be activated, but whether or not this leads to vpoDN activation and hence VPO depends on the level of pC1 activity, which is higher in virgins than in mated females. The neural computation that underlies this state-dependent sensorimotor transformation remains to be determined, pending methods for simultaneously manipulating and recording from all three cell types. One possibility is that the pC1 inputs gate the vpoEN inputs in a non-linear fashion. We did not however note any obvious spatial segregation of vpoEN and pC1 synapses onto the vpoDN dendrites, as one might expect if these inputs were processed hierarchically. Alternatively, vpoDNs might simply use a sum-to-threshold mechanism, in which the combined input from vpoENs and pC1s must exceed a certain value to elicit action potentials in vpoDNs. In this scenario, the lower level of pC1 activity after mating would necessitate a stronger vpoEN input in order to activate the vpoDNs. This model may account for the observation that wild-caught females are often multiply mated^36,37^, consistent with the prediction from evolutionary theory that a mated female would increase her reproductive fitness by re-mating if courted by a male of higher quality than her first partner. If male quality is indicated by the IPI and amplitude of his pulse song, and hence encoded in vpoEN activity, then the circuit we describe here should be capable of implementing this mating strategy.

We also note the many other, as yet uncharacterised, inputs to pC1, vpoEN, and vpoDN cells, all of which may convey additional signals that modulate female receptivity. For example, pC1 cells are reported to respond to a male pheromone^3^, which may serve to enhance the receptivity of both virgin and mated females. In this regard, it is interesting to note the persistent enhancement of vpoDN song responses upon transient activation of pC1 cells. This effect resembles the persistent state of courtship arousal induced in males by transient activation of the male pC1 counterparts^38,39^, although it lasts only seconds in females but minutes in males. The pC1 cells may respond to an initial mating in females but not males, but in both sexes may also be capable of encoding a lasting state of mating arousal induced by sensory cues that signal the general presence of potential mates. The ensuing acts of courting and mating with a specific partner, however, require a coordinated interplay of signals and responses, such as song production and VPO. These sensorimotor transformations may not be directly mediated by pC1 cells, as previously thought^40^, but nonetheless be sensitive to the arousal states they encode. The neural architecture we report here for the control of *Drosophila* female sexual receptivity may thus serve as a paradigm for understanding male sexual behaviour, and perhaps more generally for state-dependent signal processing in other behavioural decisions in flies and other species.

## Methods

### Flies

Flies were reared on standard cornmeal-agar-molasses medium except the females used for egg-laying test, which were kept on protein enriched food^41^ after mating. Flies were raised at 25 °C with relative humidity of ~50% and a 12 h/12 h light/dark cycle, unless otherwise noted.

### Split-Gal4 screening and stabilization

Split-Gal4 lines used in this study were screened and generated as described previously^4^.

### Neuron tracing in FAFB

We manually traced the neuron skeletons in a serial section transmission electron microscopy volume of the adult female *Drosophila* brain^30^, using the annotation software CATMAID^42^ (http://www.catmaid.org) as previously described^4^. We used confocal image stacks of the target cell types acquired with light microscopy as a guide to find and identify the same cell types in FAFB. We used neuroanatomical landmarks in the EM volume such as fiber tracts, cell body size and position, and neuropil boundaries to search for potential candidates of vpoDN. We looked for distinguishing features such as cell body position and tract orientation, and overall dendritic projection patterns in the confocal images. We then searched for corresponding areas of cell body position in the EM volume and followed the primary neurite emerging from the cell body as it formed fiber bundles and traversed the brain in an orientation that matched the data in the confocal images. We traced just enough of the primary and secondary neurites (backbone) of each potential candidate to compare with confocal data, and neurons that lacked prominent morphological features in the EM volume were eliminated from consideration. We identified synapses on these neurons using previously described criteria for a chemical synapse^30^. In brief, we marked instances where vpoDN was postsynaptic indicated by the presence of postsynaptic densities (PSDs) on vpoDN and by the presence of a T-bar and vesicles at an active zone in the presynaptic partner across a synaptic cleft. After the vpoDN was traced to completion and all PSDs were marked, we used CATMAID’s “reconstruction sampler” tool to randomly select upstream partners of the vpoDN which were then manually traced to identification. Using the sampler tool the reconstructed vpoDN skeleton was divided into intervals of 5000 nm. Within each interval, the sampler lists the upstream or downstream connections of the neuron that were previously marked. The sampler selects a random synapse within a given interval, for which we identified the pre-synaptic T-bar and manually traced the neuron it belonged to. All upstream partners were selected in this manner, and each was traced completely in the region of overlap with the vpoDN, and sufficiently to identify it.

### Fluorescent staining and confocal imaging

Immunofluorescence staining^4^ and fluorescent in situ hybridization^43^ (FISH) were performed as described previously. Confocal microscopy and image analysis were done as described previously (Wang et al., 2020).

### Calcium imaging

Both *ex vivo* and *in vivo* calcium imaging were performed on a customized two-photon microscope as described previously^4^. Sample preparation was as described, except that, for *in vivo* imaging, two forelegs were immobilized by applying small amounts of UV curing adhesive (Loctite 352, Henkel, Germany) to the forelegs to prevent them from touching the antennae. A loudspeaker was placed ~20 cm away from the back of the fly to play sound. In some experiment, a small amount of UV curing adhesive was used to immobilize the aristae. The songs were either recorded during fly courtship by using particle-velocity microphones (NR-23158, Knowles, IL), or synthesized in MATLAB (Mathworks, MA). Analysis of calcium imaging data was done in Fiji^44^ and MATLAB as described previously^4^.

### Electrophysiology

Whole-cell recordings were performed on CNS explants as described previously^4^.

### Behavioural assays and analysis

The flies used in behavioural assays were collected, reared, and videotaped from above as described previously^4^. Videos were taken at 30 fps with a resolution of 0.02 mm/pixel unless otherwise noted. Infrared illumination (880 nm) as well as stimulations for optogenetic activation (635 nm, 57 μW/mm^2^) or silencing (560 nm, 10 μW/mm^2^, and 635 nm, 57 μW/mm^2^) were provided from below. In experiments where a female was paired with a male, low level of constant blue light (470 nm, 0.5 μW/mm^2^) was provided for the flies to see each other.

For evaluating VPO by female flies, courtship chambers (diameter = 18 mm, height = 2 mm) were used to house single females or male-female pairs. For examining VPO at higher resolution from the ventral side, females were chilled on ice for ~30 s, and glued on a glass with ventral side facing above. Small amounts of UV curing adhesive were applied at the back of thorax and back of abdomen to minimize the movement of the fly. The field-of-view of camera was zoomed in and focused at the tip of abdomen with a resolution of 1.8 μm/pixel. Female receptivity and egg-laying by females were carried out as described previously^4^.

For annotating male behaviours around the onset of VPO by females, videos were manually analyzed offline in Fiji. The onset of female VPO was defined as the frame in which the vaginal plates open. Wing extension was defined as frames in which the male extended its wings in a singing posture. Proboscis extension was defined as frames in which the male extended its proboscis to reach female’s abdomen or genitalia. Abdomen bending was defined as frames in which the male bent its abdomen such that a line connecting the haltere and the abdominal tip came to meet at an angle of 15° or larger to the thoracic midline. Licking was defined as frames in which the male’s proboscis touched female’s genitalia. Holding was defined as frames in which the male held the female’s abdomen with two forelegs. Copulation attempt was defined as the frame in which the male’s genitalia touches the female’s genitalia.

### Statistical analysis

All egg-laying, calcium imaging and electrophysiology data were analyzed by unpaired Wilcoxon signed-rank test. All analyses were performed using R software or MATLAB.

## Acknowledgements

We thank the Janelia FlyLight, Fly Facility, Project Technical Resources, Molecular Biology, Functional Connectome, and Experimental Technology teams and Nan Chen for technical assistance, and Karel Svoboda, Kai Feng and Krystyna Keleman for comments on the manuscript. This work was funded by the Howard Hughes Medical Institute.

## Author Contributions

B.J.D., K.W., and F.W. conceived the study and wrote the manuscript. K.W. and F.W. performed all experiments and analysed the data. N. F., T.Y., C.P., and F.W. reconstructed selected neurons and synapses in the FAFB EM volume. R.P. managed the tracing team.

## Author Information

Reprints and permissions information is available at www.nature.com/reprints.

The authors declare no competing interests.

Correspondence and requests for materials should be addressed to.

---

**Extended Data Fig. 1.**
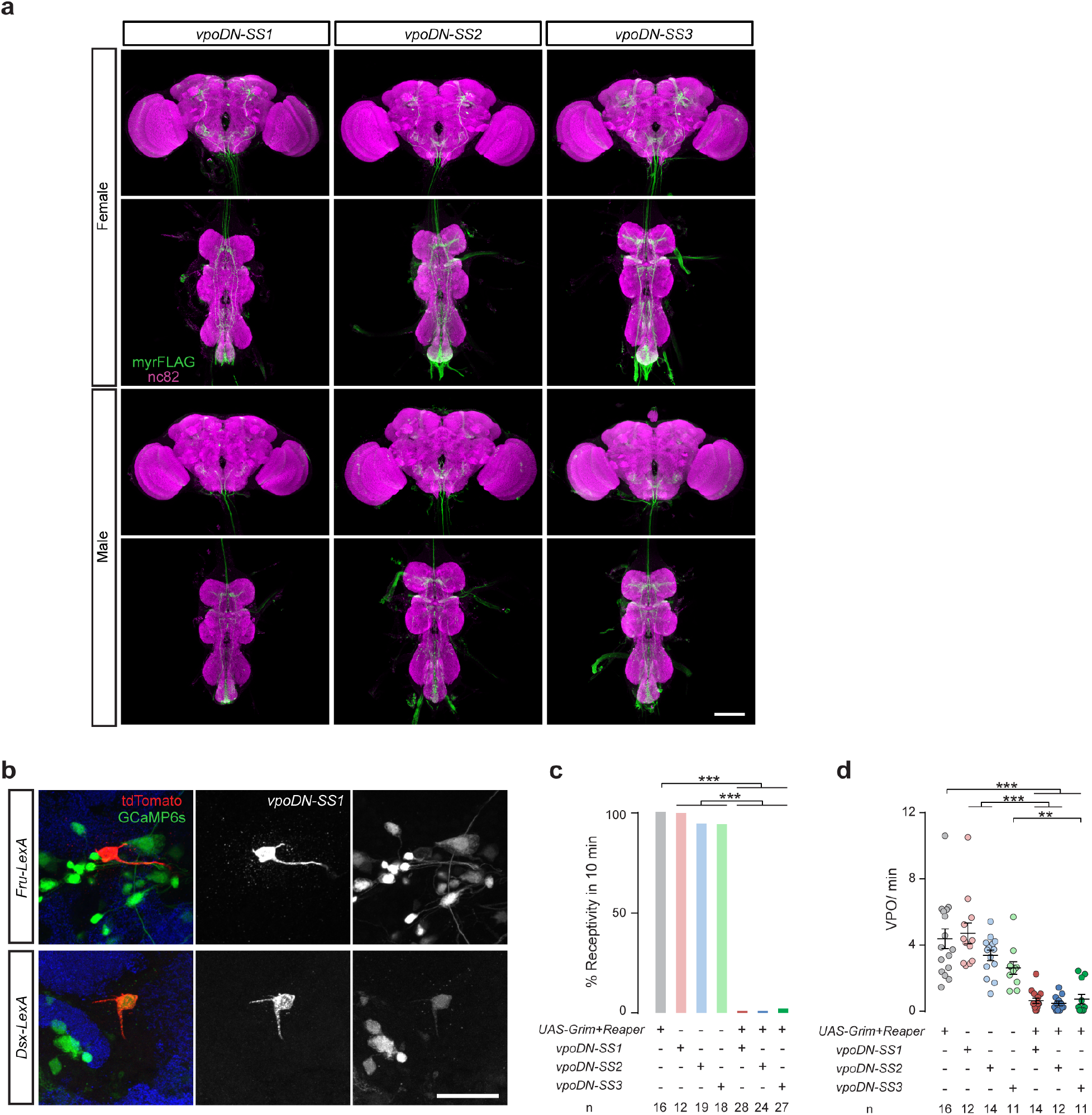
Anatomical and functional characterization of vpoDNs. **a**, Confocal images of brains and VNCs from female and male flies carrying *vpoDN-SS1*, *vpoDN-SS2*, or *vpoDN-SS3*, and *UAS-myrFLAG*, stained with anti-FLAG to reveal membranes of targeted neurons (green) and mAb nc82 to reveal all synapses (magenta). One pair of vpoDNs are labeled only in females but not in males. Scale bar: 100 μm. **b**, Confocal images of female brains showing the co-labeling of vpoDNs with *dsx-LexA* but not *fru-LexA*. Scale bar: 20 μm. **c**, **d**, Percentage of virgins copulating (**c**) and frequency of VPO (**d**) during 10 mins of courtship by a wild-type male. ***, *P* < 0.001, **, *P* < 0.01, Fisher’s exact test in **c**, Wilcoxon test in **d**. Data in **d** shown as scatter plots with mean ± s.e.m..

**Extended Data Fig. 2.**
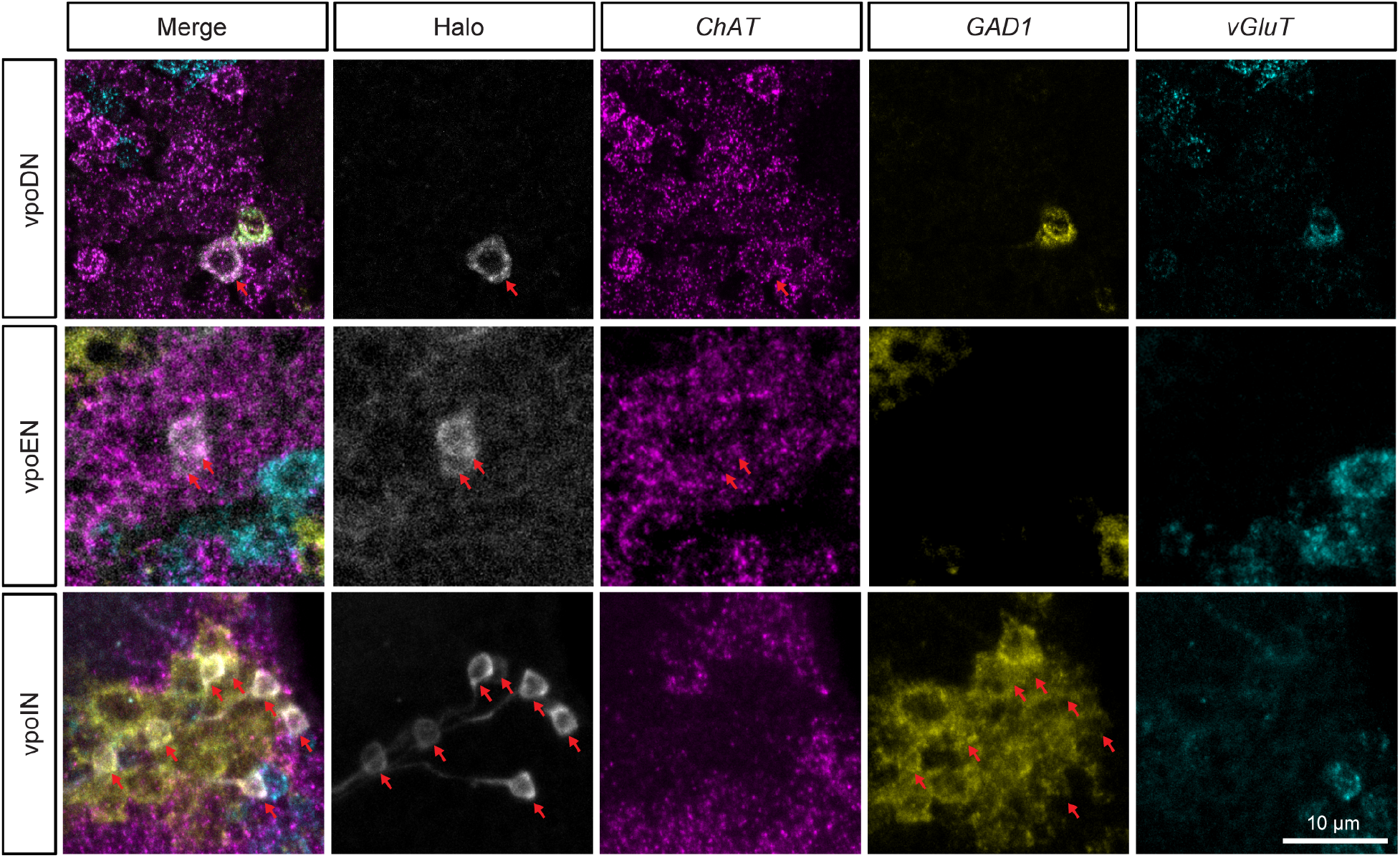
Neurotransmitter types revealed by FISH. Confocal images showing the expression of *ChAT*, *GAD1*, and *vGluT* in vpoDN, vpoEN, and vpoIN neurons in female brains. Cell bodies of interest are indicated by arrows.

**Extended Data Fig. 3.**
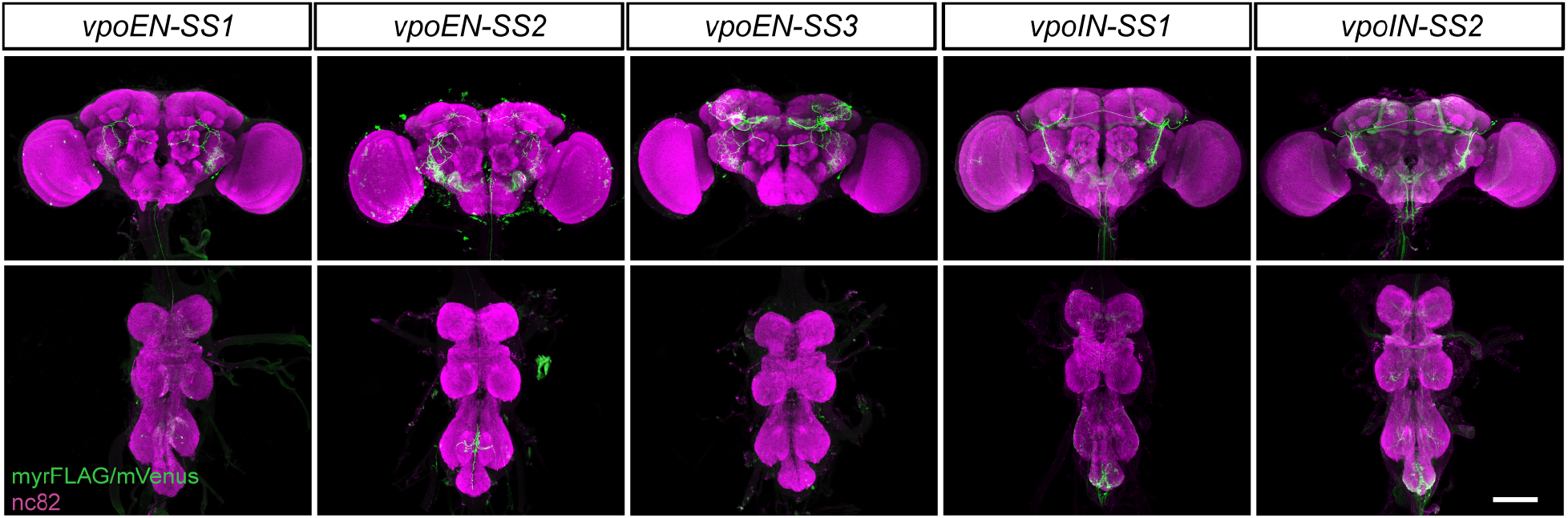
Split-Gal4 driver lines targeting vpoENs and vpoINs. Confocal images of female central nervous system carrying indicated SS driver lines and *UAS-myrFLAG* or *UAS-Chrimson-mVenus*. Samples were stained with anti-FLAG or anti-GFP (green) to reveal membranes of targeted neurons and mAb nc82 to reveal all synapses (magenta). Scale bar: 100 μm.

**Extended Data Table 1.** vpoDN inputs identified by EM reconstruction. Number of synaptic connections identified between various input neurons and the right hemisphere vpoDN (threshold 10 synapses). R and L indicate soma location in right (ipsilateral) or left (contralateral) hemisphere; N.D., soma not identified.

| Cell ID | Hemisphere | Cell type | vpoDN<br>(5094165) |
| --- | --- | --- | --- |
| 3807213 | R | pC1a | 139 |
| 3838016 | L | pC1a | 115 |
| 5787563 | R |  | 114 |
| 5496947 | R | vpoEN | 81 |
| 3794184 | R | pC1c | 77 |
| 7965359 | N.D. |  | 66 |
| 5480915 | R | vpoEN | 62 |
| 5114793 | R | vpoIN | 60 |
| 6481209 | R | vpoIN | 54 |
| 5934095 | L |  | 49 |
| 5310407 | R | vpoIN | 44 |
| 5486635 | R | vpoIN | 42 |
| 6170157 | R |  | 37 |
| 1945493 | R | vpoIN | 36 |
| 5500623 | R | vpoIN | 29 |
| 6068590 | R |  | 26 |
| 7125206 | R | vpoIN | 26 |
| 2135548 | R |  | 25 |
| 5557175 | R |  | 25 |
| 5753207 | R | vpoIN | 24 |
| 3397581 | R |  | 23 |
| 5297515 | R |  | 23 |
| 5592797 | R | vpoIN | 19 |
| 5105812 | R |  | 17 |
| 6185015 | R |  | 17 |
| 6051130 | R |  | 17 |
| 6764899 | R |  | 15 |
| 7124679 | L |  | 15 |
| 6913699 | L |  | 15 |
| 6172298 | R |  | 15 |
| 3205032 | L |  | 14 |
| 6105481 | R |  | 14 |
| 5174016 | L |  | 14 |
| 7404025 | N.D. |  | 14 |
| 3781622 | R | pC1b | 13 |
| 3778246 | R | pC1d | 13 |
| 8551397 | R |  | 13 |
| 8275720 | N.D. |  | 12 |
| 8125540 | R |  | 11 |
| 3837770 | L | pC1d | 10 |
| 5153878 | R |  | 10 |
| 5874280 | L | vpoEN | 10 |
| 5888229 | L |  | 10 |
| 7957672 | N.D. |  | 10 |
| 7979198 | R |  | 10 |

**Extended Data Table 2.**
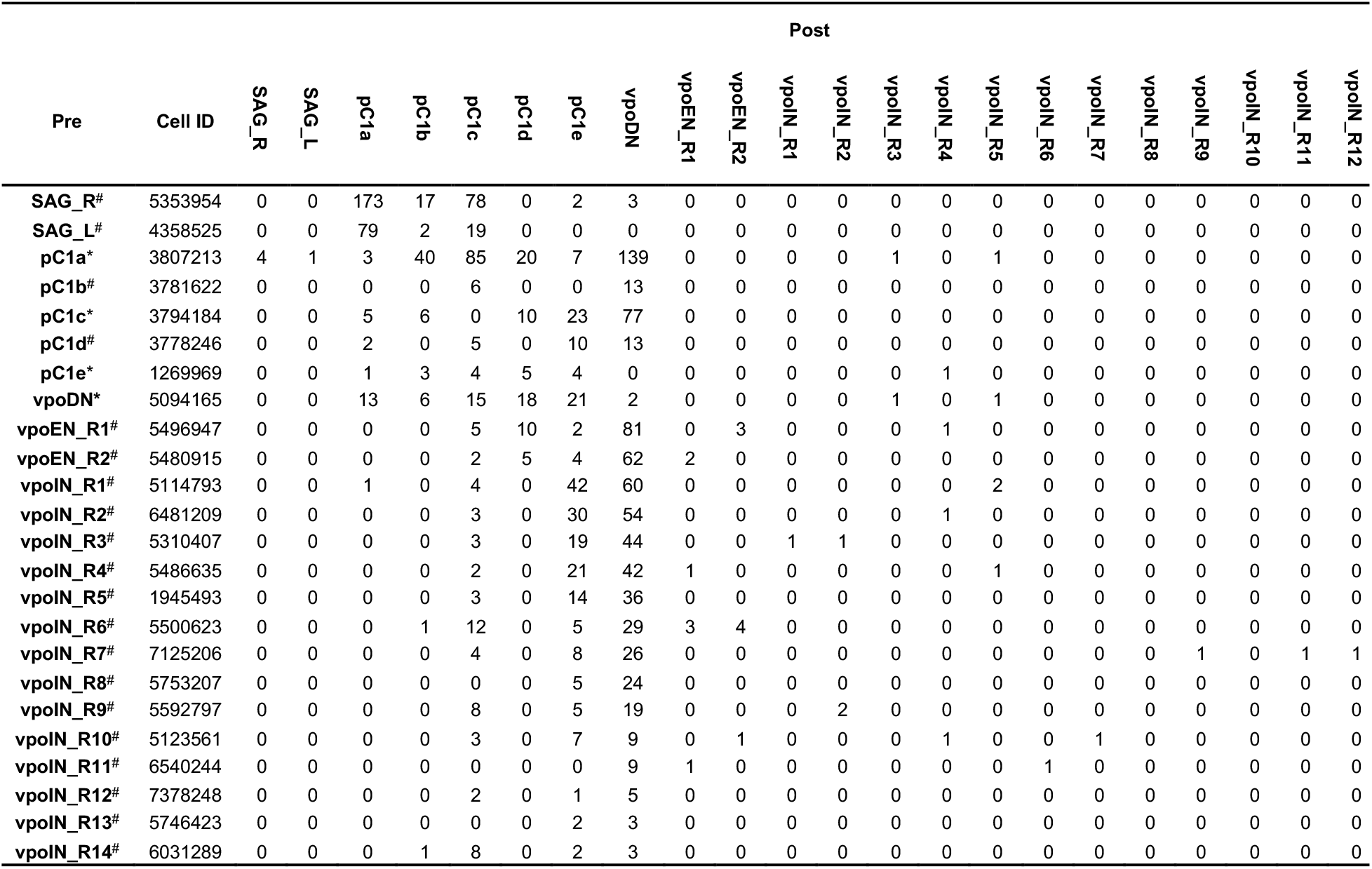
Synaptic connections identified by EM reconstruction. * fully-traced cells. ^#^ partially-traced cells. SAG_R and SAG_L indicate right and left hemisphere SAG cells, respectively. All other neurons are right hemisphere cells.

**Video 1 | Vaginal plate opening.** A virgin female fly performs VPO while being courted by a male, shown at half speed (15 fps) and replayed once. The full arena is shown on the left; a close-up of the female’s abdomen on the right.

**Video 2 | Optogenetic activation of vpoDNs elicits vaginal plate opening.** A montage of video clips of 4 virgin *vpoDN-SS1 UAS-Chrimson* females upon 3 s of continuous 625 nm illumination at 200 μW/mm^2^), shown at normal speed (30 fps).

**Video 3 | vpoDN reconstructed in female brain EM volume.** A single vpoDN cells (red) reconstructed in the right hemisphere of the FAFB EM volume.

